# A genome-wide interactome of DNA-associated proteins in the human liver

**DOI:** 10.1101/111385

**Authors:** Ryne C. Ramaker, Daniel Savic, Andrew A. Hardigan, Kimberly Newberry, Gregory M. Cooper, Richard M. Myers, Sara J. Cooper

## Abstract

Large-scale efforts like the Encyclopedia of DNA Elements (ENCODE) Project have made tremendous progress in cataloging the genomic binding patterns of DNA-associated proteins (DAPs), such as transcription factors (TFs). However most chromatin immunoprecipitation-sequencing (ChIP-seq) analyses have focused on a few immortalized cell lines whose activities and physiology deviate in important ways from endogenous cells and tissues. Consequently, binding data from primary human tissue are essential to improving our understanding of in vivo gene regulation. Here we analyze ChIP-seq data for 20 DAPs assayed in two healthy human liver tissue samples, identifying more than 450,000 binding sites. We integrated binding data with transcriptome and phased whole genome data to investigate allelic DAP interactions and the impact of heterozygous sequence variation on the expression of neighboring genes. We find our tissue-based dataset demonstrates binding patterns more consistent with liver biology than cell lines, and describe uses of these data to better prioritize impactful non-coding variation. Collectively, our rich dataset offers novel insights into genome function in healthy liver tissue and provides a valuable research resource for assessing disease-related disruptions.

## Introduction

Complex gene regulatory networks underlie key aspects of human development, tissue physiology, and cell fate determination (Karlebach and Shamir 2008; Spitz and Furlong 2012). These gene expression programs are coordinated by DNA-associated proteins (DAPs), especially sequence-specific transcription factors (TFs), which bind to promoters, enhancers, silencers, insulators and other c/s-regulatory elements (Spitz and Furlong 2012). Owing to their fundamental biological importance, disease can result from disruption or alteration of transacting DAPs or the c/s-regulatory elements to which they bind (Sur and Taipale 2016; Khurana et al. 2016). Accordingly, the interactions of DAPs and regulatory sequences has been investigated extensively (Birney et al. 2007; Bernstein et al. 2012; Gerstein et al. 2012; Andersson et al. 2014). These studies have been greatly aided by high-throughput sequencing technologies to map genome-wide binding patterns of DAPs, in particular via chromatin immunoprecipitation sequencing (ChIP-seq) (Johnson et al. 2007; Robertson et al. 2007).

The vast majority of genome-wide DAP binding maps, such as those from the Encyclopedia of DNA Elements (ENCODE) Project (https://www.encodeproject.org), are generated in a small number of mostly tumor-derived cell lines. Such data are clearly useful for understanding basic genome-wide protein-DNA interactions and allow for a variety of experimental perturbations (Savic et al. 2015, 2016; Reddy et al. 2012; Gertz et al. 2012a). However, these in vitro systems are likely to be limited in the extent to which they recapitulate *in vivo* tissue environments, especially for non-cancerous tissues (Ertel et al. 2006; Sandberg and Ernberg 2005).

The generation of genome-wide, DAP binding patterns in healthy human tissue is essential to improving our understanding of transcriptional control within a physiological context. Comprehensive analyses of genome-wide DAP binding sites across a variety of different tissue types (Savic et al. 2013) and individuals can serve as a critical complement to existing cell line-based catalogues. In support of these ideas, tissue-based studies have successfully defined *cis-* regulatory elements critical for tissue differentiation and disease progression (Blow et al. 2010; Visel et al. 2013). Other investigations have provided evolutionary insights (Schmidt et al. 2010; Stefflova et al. 2013) and have highlighted the impact of non-coding, regulatory variation on drug response (Soccio et al. 2015). In addition, a comprehensive investigation of regulatory element usage across murine tissues and developmental time points has further illustrated the plasticity and spatiotemporal specificity of gene regulatory architecture (Nord et al. 2013).

We generated ChIP-seq data for twenty DAPs in human liver tissue samples from two donors, one a young female (4 years of age) and the other an adult male (31 years of age), (Supplemental Table 1 and 2). These donors did not have any chronic or acute illnesses that might compromise their liver function and both succumbed to fatal accidents that spared their livers from observable damage. We selected DAPs based on the availability of a suitable ChIP-seq grade antibody and their expression levels in the liver, ultimately assaying 17 sequence-specific TFs, CTCF and RAD21, two DAPs involved in maintaining chromatin structure, and RNA polymerase II (RNAP2), which is directly involved in transcription. We also generated RNA-sequencing (RNA-seq) and whole genome data from the same samples. This dataset provides the most comprehensive genomic characterization of DAPs in a healthy tissue to date and is freely available (https://www.encodeproject.org).

Here we present a broad analysis of these data, including an assessment of DAP interactions and a comparison of DAP occupancy between donor tissues and a widely studied liver cancer derived cell line (HepG2). We also analyzed genomic regions previously associated with liver biology or disrupted in liver cancer and describe an approach for using our DAP occupancy data to prioritize impactful non-coding variation. Binding data was integrated with previous histone modification data derived from liver tissue by the Epigenome Roadmap Consortium (http://www.roadmapepigenomics.org), tissue expression and expression-QTL data from the Genotype-Tissue Expression Project (http://www.gtexportal.org/), and hepatocellular carcinoma whole genome somatic mutation data from the International Cancer Genome Consortium (https://dcc.icgc.org). Our results provide compelling insights into human liver transcription and provide a valuable resource for future analyses of both mechanisms of gene regulation and relevance of non-coding variation to disease.

## Results

### DAPs display extensive co-localization

Liver tissue samples were acquired from a 4 year old female donor and a 31 year old male donor with consent from both donor families. We performed pairs of replicate ChIP-seq assays for each of the 20 DAPs (Supplemental Table 1) in both tissue samples, resulting in a total of 80 independent ChIP-seq experiments. ChIP-seq experiments were conducted in accordance with ENCODE guidelines (Landt and Marinov 2012). All replicate pairs were strongly correlated and canonical motif enrichment was detected for all sequence-specific TFs (Supplemental Table 3, Supplemental Material). We identified between 909 and 60,597 binding events for each DAP and tissue sample (Supplemental Table 1), and identified more than 450,000 binding sites, spread over 150,000 unique genomic locations, across all DAPs in both livers.

We first sought to examine relationships between DAP binding profiles in each liver that might suggest interactions between factors. Hierarchal clustering of normalized ChIP-seq read count correlations within the union of all binding sites revealed strong correlations between many pairs of DAPs in both tissues (median rho of 0.718 and 0.642, and max rho of 0.889 and 0.863, in the male adult and female child livers respectively), with no factors negatively correlated (Figure 1A-B). We further identified largely consistent binding patterns between the two tissue samples despite age and sex differences, with ~75% of all DAP binding sites shared between the two donors (Figure 1C). We observed stronger binding similarity between maps of a given DAP from the two samples than between a given DAP and other factors in the same sample (Wilcox *P*<0.0001, Supplemental Figure 1-2). Consistent with their roles in maintaining genome insulation and three-dimensional genome structure (Merkenschlager and Nora 2016), RAD21 and CTCF displayed the most distinctive binding patterns and clustered separately from all other DAPs in both tissues.

**Figure 1.**
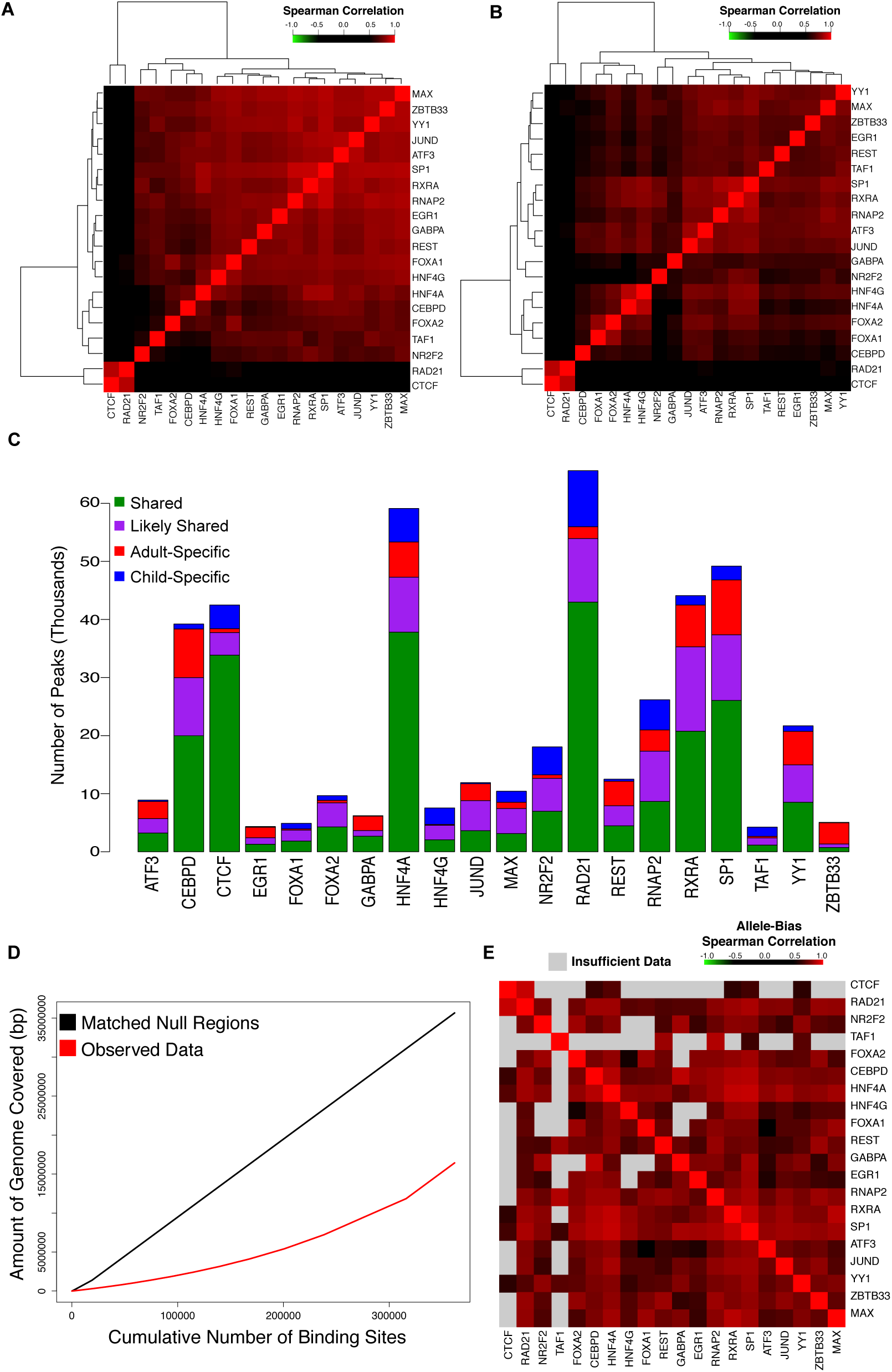
(A and B) Heatmaps of spearman correlation matrix of normalized DAP binding intensities at all observed binding sites in the adult (A) and child (B) liver respectively. (C) Stacked bar plot displaying the number of peaks for each TF. Peaks are broken in to those that are shared among both replicates of both donors (shared, green), are shared between a donor and one replicate of the other donor (likely shared, purple), are specific to the adult donor (adult-specific, red), and are specific to the child donor (child-specific, blue). (D) Cumulative number of base pairs covered per binding site included in adult liver observed data (red) and null regions (black) matched for length, GC content, and repeat content. (E) Heatmap of a pair-wise Spearman correlation matrix, ordered identically to (1A) indicating the correlation of allele bias at shared peaks from the adult donor that overlap a heterozygous SNP for each pair of factors. The color of each panel indicates the strength of the correlation with gray indicating less than 25 peaks passed the quality filters for allele bias analysis for a given pair.

To provide context to the degree of interaction between factors in the adult male liver, we used a previously described method to randomly sample genomic regions with length, GC content, and repetitive sequence content matched to that of the observed binding sites (Fletez-Brant et al. 2013). Compared to these randomly sampled regions, actual DAP binding sites covered ~50% fewer bases (Figure 1D), indicating that observed overlap rates are far above random expectation. Binding sites of FOXA1, a previously described pioneer factor (Zaret and Carroll 2011), had the highest mean number of sites overlapping with sites bound by other factors and the degree of co-localization at FOXA1 binding sites differed dramatically from non-FOXA1 bound sites (Supplemental Figure 3). We next assessed coordination in allele-specific binding among DAPs in the adult liver tissue. Using a previously described method (DeSantiago et al. 2016), we assessed the degree of allele bias, measured as the fraction of ChIP-seq reads containing the human genome (hg19) reference sequence, for each DAP at all heterozygous single nucleotide variants (SNVs) that overlapped with an adult DAP binding site. Correlation analyses of the degree of allele bias at each SNV between all possible pairs of DAPs revealed that factors also tended to cluster on the same allele, indicative of cooperative binding to the same chromosome (Figure 1E, Supplemental Table 4).

The relative importance of regions with varying degrees of DAP interaction was assessed by examining how evolutionary conservation and chromatin modifications relate to the number of factors bound at a site. We found that sites with higher numbers of DAPs tended to be more strongly conserved across the mammalian phylogeny (rho = 0.960, *P* = 7.08×10^−6^, Supplemental Figure 4A). These sites were also more heavily enriched for the activating histone 3 acetylation marks on lysine 9 (H3K9ac) and lysine 27 (H3K27ac) and relatively depleted for the repressive histone 3 methylation marks on lysine 9 (H3K9me3) and lysine 27 (H3K27me3) (Supplemental Figure 4B-E). Together, these data show that DAPs co-localize extensively, as previously demonstrated in cell line analyses (Yan et al. 2013), that they often show equivalent allele bias, and that sites of high interaction tend to be more highly evolutionarily conserved and more strongly associated with active chromatin marks.

### DAP binding recapitulates known liver expression programs

We next integrated DAP binding profiles with gene annotations in an attempt to infer DAP regulatory networks and assess the relationship between DAP occupancy and gene expression level. We found that 30% of all ENSEMBL (GRCh37_E75) annotated genes harbored at least one binding site within 1kb of their transcriptional start sites (TSSs), and more than 7% of genes harbored binding events for more than 6 different DAPs in both adult and child (Figure 2A, Supplemental Figure 5A). To identify correlations between gene expression and binding site maps, we performed quadruplicate RNA sequencing (RNA-seq) experiments on independent samples from each donor tissue. As expected, RNAP2 promoter binding within 1kb of a gene’s TSS was strongly associated with expression in both livers (Wilcoxon *P*<0.0001, mean TPM without RNAP2 = 22.3, and mean TPM with RNAP2 = 121.8). Gene expression level was also strongly correlated with the number of factors binding within 1kb of their TSS (rho=0.533 for adult and rho=0.526 for child, Figure 2B, Supplemental Figure 5B), however this effect diminished rapidly as the distance to TSS threshold expanded beyond 1kb (Figure 2C).

**Figure 2.**
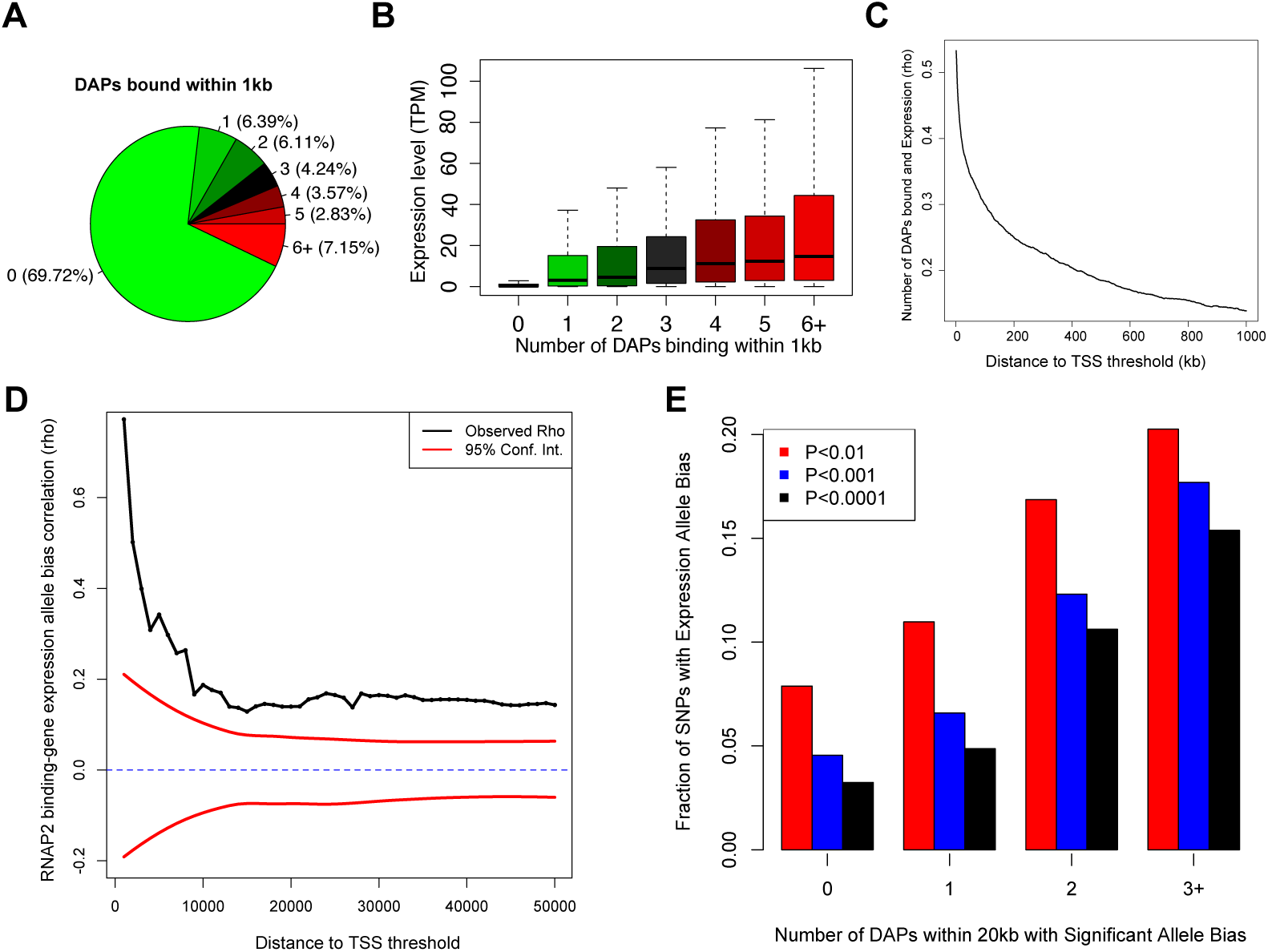
(A) Pie chart representing the percent of genes containing a specified number of bound DAPs within 1 Kb of their TSS. Color scale reflects the number of neighboring DAPs required to be included in each slice. (B) Expression level of genes binned by the number of DAPs bound within 1Kb of their TSS. Data shown for the adult donor, child data shown in Supplemental Figure 5. (C) Correlation between expression level of genes and the number of factors bound (as described in Figure 2B above) for a range of distance to TSS thresholds. (D) Correlation between RNAP2 binding and neighboring gene expression allele bias over a range of distance thresholds. Ninety five percent confidence (red) intervals calculated by randomly shuffling all SNP pairs that met a distance threshold 100 times and computing a null correlation distribution. (E) Bar plot showing the fraction of expressed SNPs with significant allele bias with varying numbers of neighboring DAPs exhibiting allele bias within 20kb. Significance thresholds of binomial test P<0.01 (red), P<0.001 (blue), and P<0.0001 (black).

Using phased whole genome sequencing data, we assessed how allele bias in DAP binding correlated with neighboring allele bias in gene expression. We found bias in RNAP2 occupancy within 1 kb of a gene’s TSS was strongly correlated (rho=0.750) with expression bias and the strength of this correlation dropped precipitously as the distance between binding site and TSS was expanded (Figure 2D). A similar pattern was observed with the other assayed DAPs, although the strength of the correlations were largely reduced relative to that observed for RNAP2 and repressive factors such as REST and NR2F2 exhibited correlations in the opposite direction (Supplemental Figure 6-7, Supplemental Table 5). We also found the number of neighboring DAPs with significant allele bias was associated with the likelihood of observing significant allele bias in gene expression (Figure 2E).

To define the gene regulatory networks associated with each DAP, we compared the distribution of the distances from the nearest DAP site to the TSSs of genes in every Reactome (http://www.reactome.org) pathway to that of the background transcriptome using a Kolgomorov-Smirnov (KS) test (Figure 3A, Supplemental Figure 8, Supplemental Table 6). This analysis revealed significant pathway enrichments for nearly all DAPs (median KS test *P*<0.05), including pathways known to be active in liver tissue such as lipid and carbohydrate metabolism, drug metabolism and complement activation. We also identified a second cluster of genetic pathways involved in regulating stem-cell state and cell division that was largely restricted to SP1, YY1, and GABPA binding events. An extreme example is the Hedgehog ‘on’ state (REACT_268718), a pathway that acts as an important regulator of animal development and differentiation (Ingham et al. 2011). SP1, YY1, and GABPA were all bound within 1kb of the TSSs of nearly 50% of the 60 genes within this pathway. By comparison, a distance threshold of 1Mb is required to achieve a similar level of occupancy for any other DAP (Figure 3B). SP1, YY1 and GABPA have been previously described as interacting partners (Galvagni et al. 2001; Rosmarin et al. 2004) and these pathway enrichments are consistent with previous studies implicating GABPA in controlling stem cell maintenance and differentiation (Yu et al. 2016).

**Figure 3.**
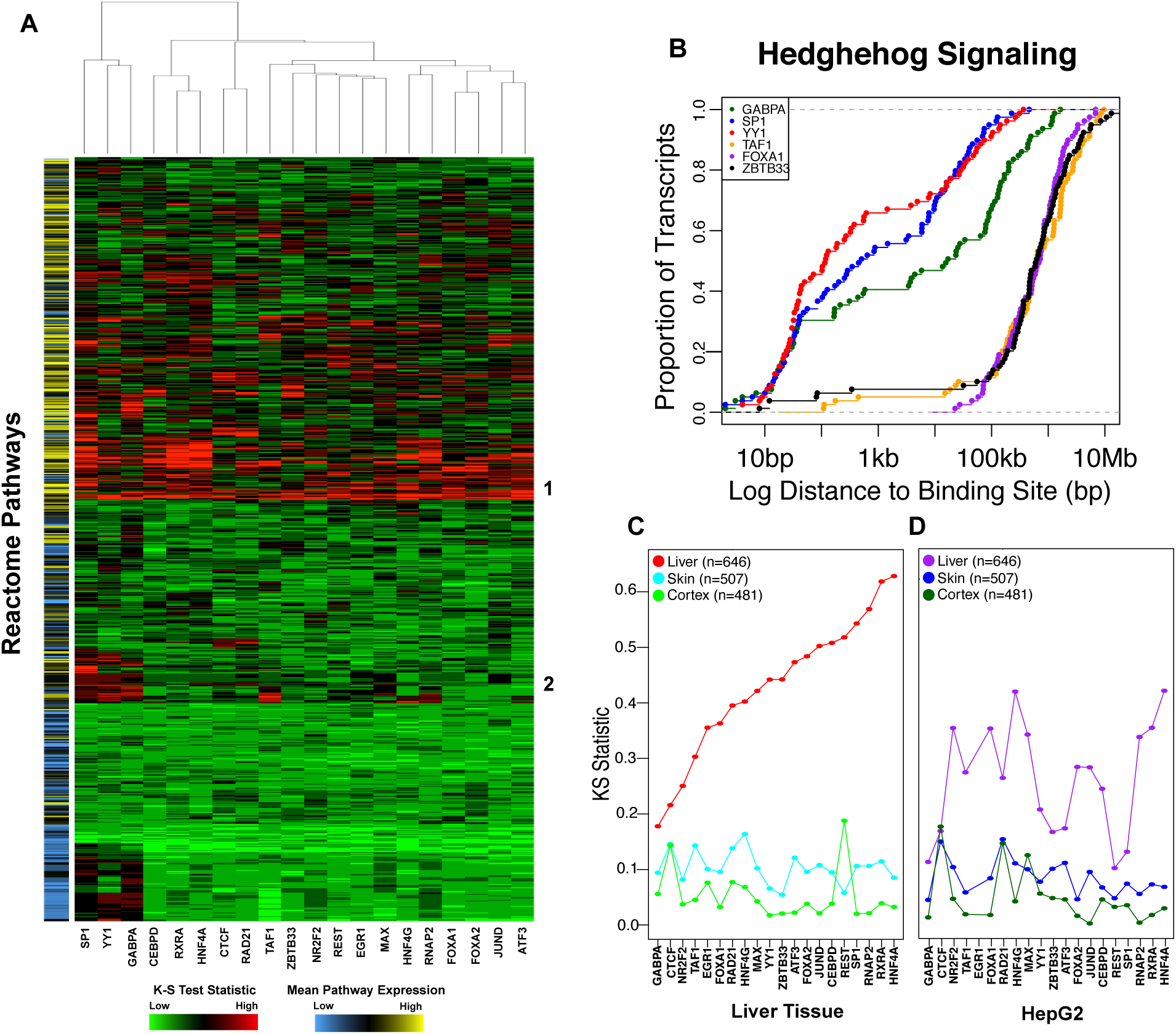
(A) Heatmap of KS-test statistic comparing distances of TSSs in a pathway to the nearest binding site for a given DAP compared to the background transcriptome for each Reactome pathway and each DAP. Concordant sites between both livers were used as input to this analysis. The color bar on the left indicates the mean expression level of genes within a pathway. (B) Representative private pathway cumulative distribution function plot demonstrating enrichment for proximal GABPA (blue), SP1 (green), and YY1 (red) binding. (C and D) Dots represent KS-statistic of enrichment for proximal binding of each factor to liver (red/purple), skin (blue) and cortex (green) -specific genes in adult liver tissue (C) and HepG2 cells (D). Data from the child donor is similar to that of the adult donor (Supplemental Figure 10).

We next analyzed promoter-proximal DAP binding to tissue-specific genes based on expression data from the Genotype-Tissue Expression Project (GTEx) (The GTEx Consortium 2015). We defined “liver-specific” genes as those with a mean RPKM, across all GTEx liver tissues, of at least two and at least five-fold higher than the mean RPKM in all other non-liver tissues. HNF4A and RXRA were the top TFs in terms of enrichment for binding near the TSSs of liver-specific genes (Figure 3C, Supplemental Figure 9), underscoring their importance in regulating liver-specific functions (DeLaForest et al. 2011; Martinez-Jimenez et al. 2010; Li et al. 2015). Supporting the tissue specificity of DAP binding events, skin and cortex-specific genes exhibited a much lower enrichment for proximal DAP binding compared to liver-specific genes. Notably, a similar analysis of ChIP-seq data generated by our group in HepG2 cells for nearly all DAPs examined here revealed enrichment for promoter-proximal binding to liver-specific genes (Figure 3D). However, aggregated across all 20 DAPs, proximal binding enrichment was significantly more pronounced in liver tissue (Paired Wilcoxon *P*=1.29×10^−4^). Only NR2F2 and HNF4G exhibited even nominally greater proximal binding enrichment in HepG2 cells than in liver tissue (Figure 3D). These results were robust to different thresholds used define tissue-specificity (Supplemental Figure 10).

### DAP binding sites are enriched for expression-QTL SNPs

We hypothesized that, by virtue of marking regulatory elements active in human liver, DAP binding sites would be enriched for expression-QTL (eQTL) SNPs previously catalogued in healthy human liver tissues by the GTEx Project. We compared the number of significant eQTL SNPs overlapping binding sites for a given DAP to 1,000 randomly sampled sets of SNPs that passed GTEx filtering and were matched for distance to nearest TSS and minor allele frequency. This analysis revealed significant enrichment (FDR<0.05) for 17 of the 20 assayed DAPs (Figure 4A, Supplemental Table 7A); JUND, FOXA1, and ATF3 were also enriched but at higher FDRs. For most DAPs, this enrichment was specific to liver eQTL SNPs. Repeating the analysis on three tissues (uterus, vagina, anterior cingulate cortex) with less than 35% eQTL SNPs shared with liver revealed significantly less overlap (Supplemental Table 7B). To examine tissue specificity more comprehensively, we assessed DAP binding site overlap with tissue-specific eQTLs, defined as those eQTLs with a FDR corrected *P* < 0.05 in only a single tissue. Bound sites of 11 of the 20 assayed DAPs overlapped with liver-specific eQTL SNPs more frequently than with eQTL SNPs specific to other tissues (Supplemental Figure 11 and Supplemental Table 8).

**Figure 4.**
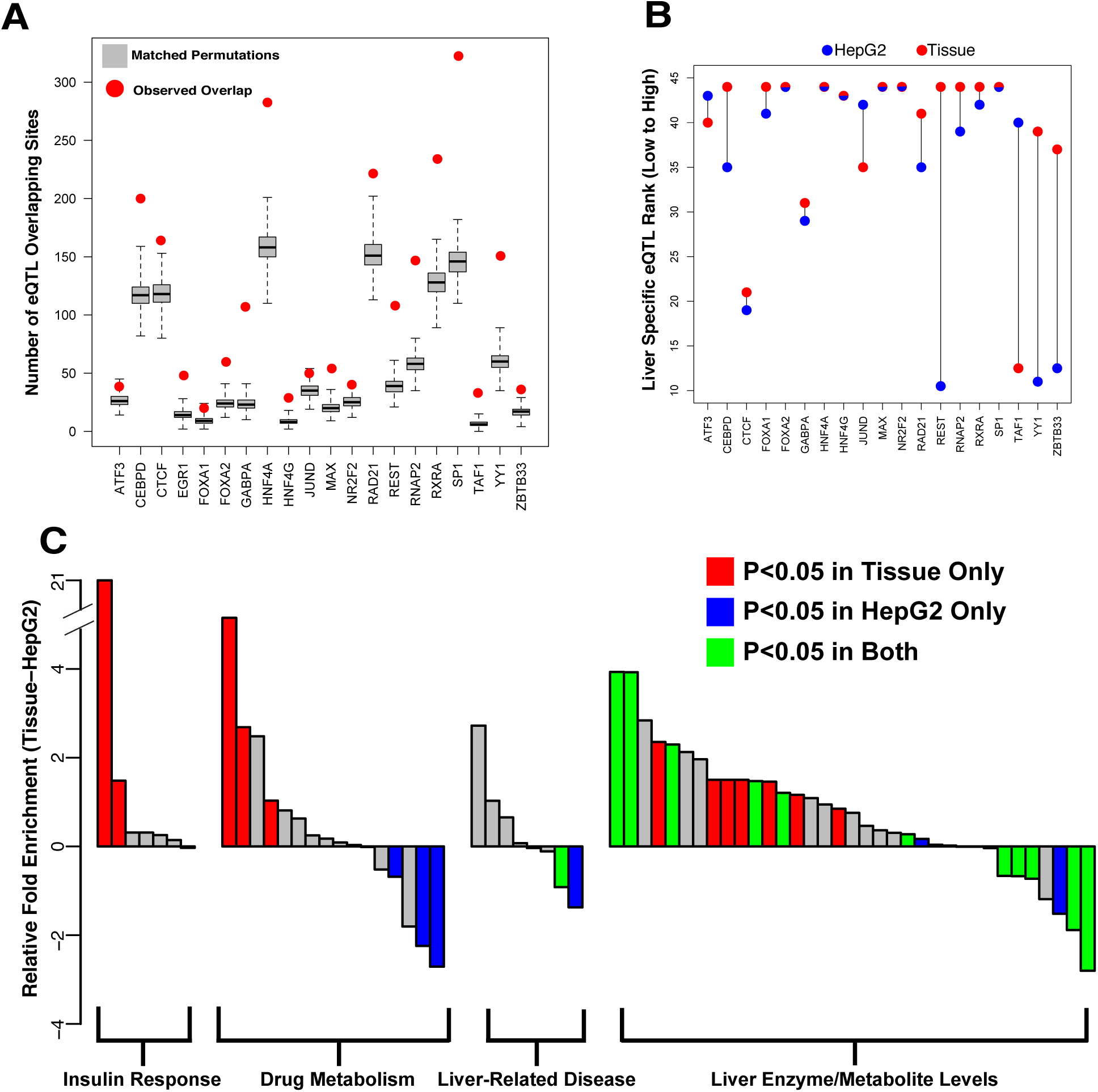
(A) Red dot indicates the number of eQTLs falling within a DAP binding site relative to the gray boxplots, which represent 1000 randomly sampled null SNPs matched for distance to TSS and minor allele frequency. (B) Relative rank of liver-specific eQTL compared to all GTEx tissue specific eQTLs in DAP binding sites assayed in HepG2 cells (blue) and adult liver tissue (red). (C) Difference in enrichment (delta Fisher’s exact test odds ratio) for SNPs associated liver-related phenotypes between binding sites for 19 common DAPs assayed in HepG2 cells and liver tissue. GWAS terms represented by bars are provided in Supplemental Table 9A.

In HepG2 cells, we observed a similarly strong enrichment for liver eQTL SNP overlap for all factors except SP1 (Supplemental Figure 12A, Supplemental Table 7A). However, this enrichment was much less specific to liver eQTL SNPs as we observed a strong enrichment in uterine, vaginal, and anterior cingulate cortex eQTLs relative to the tissue-derived DAP binding (Supplemental Table 7C). Moreover, we observed a reduction in the level of enrichment for liver-specific eQTLs relative to non-liver tissue-specific eQTLs in 10 out of 19 DAPs that were assayed in HepG2 cells, suggesting that, at least for some DAPs, tissue ChIP-seq data better identify regions important for regulating tissue-specific gene expression (Figure 4B, Supplemental Figure 12B).

We next compared HepG2 and liver tissue ChIP-seq binding site enrichment for overlap with SNPs previously associated with a liver-related phenotype in the NHGRI-GRASP genome wide association study (GWAS) catalog (https://grasp.nhlbi.nih.gov/Overview.aspx). We found greater enrichment (Paired Wilcoxon *P*=1.8×10^−3^J in liver tissue binding sites for a majority of GWAS terms (45 of 66), including insulin-like growth factor levels and response to statins (Figure 4C, Supplemental Table 9). However, HepG2 binding sites also showed enrichment for several terms, such as liver cancer risk and HDL cholesterol levels, at a higher level than that observed in liver tissue.

### DAP binding analyses help prioritize impactful non-coding variation

One of the promises of high-throughput cataloging of DAP binding is its utility in prioritizing noncoding variation capable of disrupting regulatory elements. A challenge associated with using ChIP-seq data for this purpose is that DAP occupancy peaks are often broad, and it is unclear what proportion of sequence variation within a ChIP-seq peak is likely to significantly affect DAP binding dynamics. We therefore assessed the degree of mammalian evolutionary sequence conservation within ChIP-seq peaks using Genomic Evolutionary Rate Profiling (GERP) scores (Cooper et al. 2005). The mean GERP-RS score of each protein’s binding sites was significantly greater than the genome-wide average but lower than that of protein-coding exons (Supplemental Figure 13). While these data likely reflect reduced non-coding relative to coding constraints, they may also be due to the fact that ChIP-seq-defined DAP binding sites do not have the resolution to identify the most critical nucleotides for DAP binding, such as TF motifs, which are often more highly conserved than surrounding sequences (Bernstein et al. 2012; Timothy et al. 2012). However, recent sequence-based machine learning and allele-specific binding data have also indicated that the most important sequence elements for DAP binding may not necessarily reside solely in the canonical DNA sequence motif (Deplancke et al. 2016; Timothy et al. 2012; Tehranchi et al. 2016).

Consequently, to systematically examine evolutionary conservation at base pairs most critical for DAP binding, we applied a previously described method (Lee 2016), to train 10mer-based support vector machines (SVMs) capable of distinguishing binding sites identified in DAP ChlP-seq experiments from genomic loci matched for GC and repeat content, and without evidence of DAP binding (Figure 5A). These SVMs were successful in predicting DAP binding for all factors with a mean Receiver-Operator Characteristic Area Under the Curve (ROC-AUC) of 0.928 and Precision Recall Area Under the Curve (PR-AUC) of 0.702 (Supplemental Figures 14A and 14B). A subset of 10mers, each occurring in a small percentage of total binding sites, were most predictive of DAP binding (Supplemental Figure 14C). We next computed a “delta” binding score for all possible point mutations within DAP binding sites, defined as the mean decrease in our SVMs’ classifier value for the mutant relative to the reference sequence. This strategy is similar to a previously developed approach, ‘deltaSVM’, that focuses on more local disruptions of 10mer feature weights (Lee et al. 2015). Notably, bases with the most negative delta binding score tended to be the most highly conserved for most DAPs (Supplemental Table 10). DAPs with relatively low mean binding site GERP scores, such as GABPA and CTCF, harbored high levels of conservation at their most putatively vulnerable nucleotide position (Figure 5B). We also observed a modest, but significant, correlation between delta binding scores and the observed degree of allele bias (FDR<0.05) in binding sites for 13/20 DAPs (Figures 5C, Supplemental Figure 15, Supplemental Table 11), further supporting our confidence in predicting putatively impactful variation at DAP binding sites.

**Figure 5.**
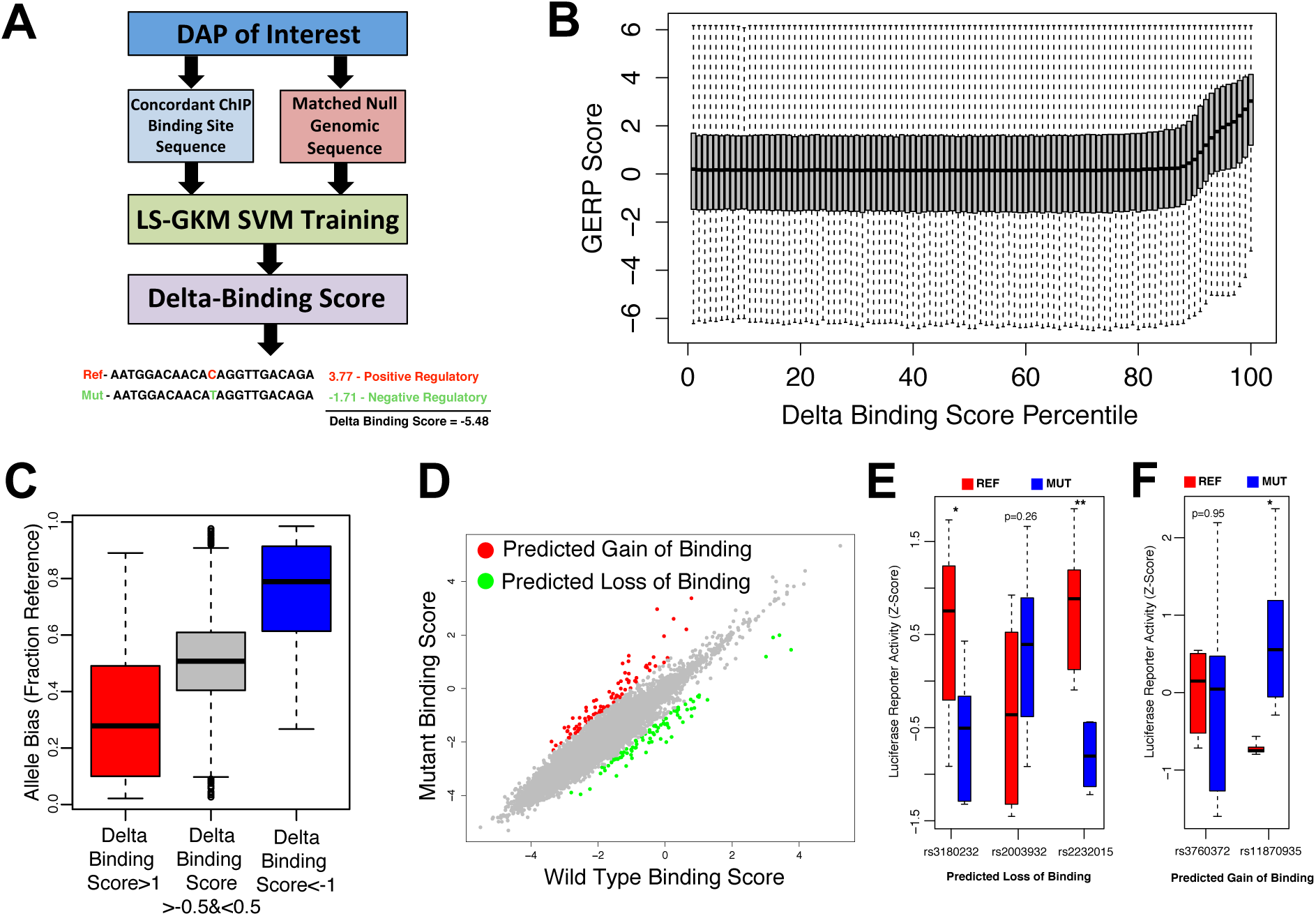
(A) Diagram representing the pipeline for generating SVMs capable of distinguishing DAP binding sites from matched null regions and scoring the predicted impact of each mutation on each DAP. (B) Boxplots representing the GERP scores at each delta binding score percentile (higher percentile indicates increased predicted disruption of binding) from *in silico* mutations for CTCF. (C) Boxplots of CTCF binding site-overlapping heterozygous SNPs predicted to be in the top ~1% for decreasing binding affinity (red), to be in the top ~1% for increasing binding affinity (blue), and to have no significant impact on binding affinity (black). Y-axis indicates the fraction of ChIP reads mapping to the reference allele (D) SVM scores for reference and alternate GTEx liver eQTL SNP alleles for CTCF. Red dots indicate SNPs that hold a positive delta binding score in the top 0.1 percentile. Blue dots indicate SNPs that hold a negative deltabinding score that falls in the bottom 0.1 percentile of all scores. (E and F) Boxplots representing the luciferase activity of reference (red) and mutant (blue) sequence in eQTL SNPs predicted to disrupt observed DAP binding (E) and induce aberrant TF binding (F). * indicates a two-tailed t-test *P*<0.05 and ** indicates a *P*<0.005.

We subsequently searched among all GTEx liver eQTL SNPs for those most likely to alter a DAP binding site (Figure 5D). Because identifying common sequence variation with functional significance is challenging for eQTL analyses and genome-wide association studies (GWASs) (Edwards et al. 2013), weighting SNPs based on their likelihood to impact DAP binding could be a useful approach for prioritizing follow-up analyses of SNP associations. Considering the top 0.1% of DAP-disruptive eQTL SNPs, we noted several that had been associated with one or more relevant phenotypes by querying the NHLBI-GRASP GWAS catalog (Supplemental Tables 12 and 13). Supporting the biological relevance of these analyses, the top 0.1% of our putative DAP-disrupting SNPs were significantly enriched (Fisher’s *P*<0.05) compared to all significant liver eQTL SNPs for various liver related GWAS catalog terms, including lipid level measurements, alcohol dependence, vitamin D levels, and levels of multiple enzymes produced by the liver (Supplemental Figure 16). To validate DAP binding disruptions, we selected two putative SNPs predicted to induce cryptic binding, as well as three SNPs predicted to disrupt binding, to test with a luciferase reporter assay in HepG2 cells (Supplemental Table 14, Figure 5E-F). Predicted disruptions were confirmed for three out of the five SNPs tested (two loss of binding and one gain of binding) confirming our approaches ability to identify SNPs with potential regulatory impact.

### Disruption of DAP activity in hepatocellular carcinoma

Our catalog of DAP binding provides an opportunity to assess cancer-related disruptions of gene regulation in healthy tissue. An examination of the degree of adult DAP binding enrichment near genes differentially expressed between The Cancer Genome Atlas project (TCGA, https://cancergenome.nih.gov) hepatocellular carcinoma tumor and adjacent normal tissue (DESeq2, FDR<0.001) revealed a general trend for preferential DAP localization near genes down-regulated in liver cancer (Figure 6A). This is consistent with a model wherein the majority of DAP-associated regulatory elements function to maintain cell identity and differentiation, both of which are widely disrupted during tumorigenesis (Sur and Taipale 2016). GABPA and SP1 stood out as outliers with strong enrichment (FDR<0.05) for binding in proximity to genes up-regulated in tumor tissue. This may be due to the importance of GABPA and SP1 proteins for regulating stem cell state and cell division as described in the Reactome pathway enrichment analysis performed above (see Figure 3A).

**Figure 6.**
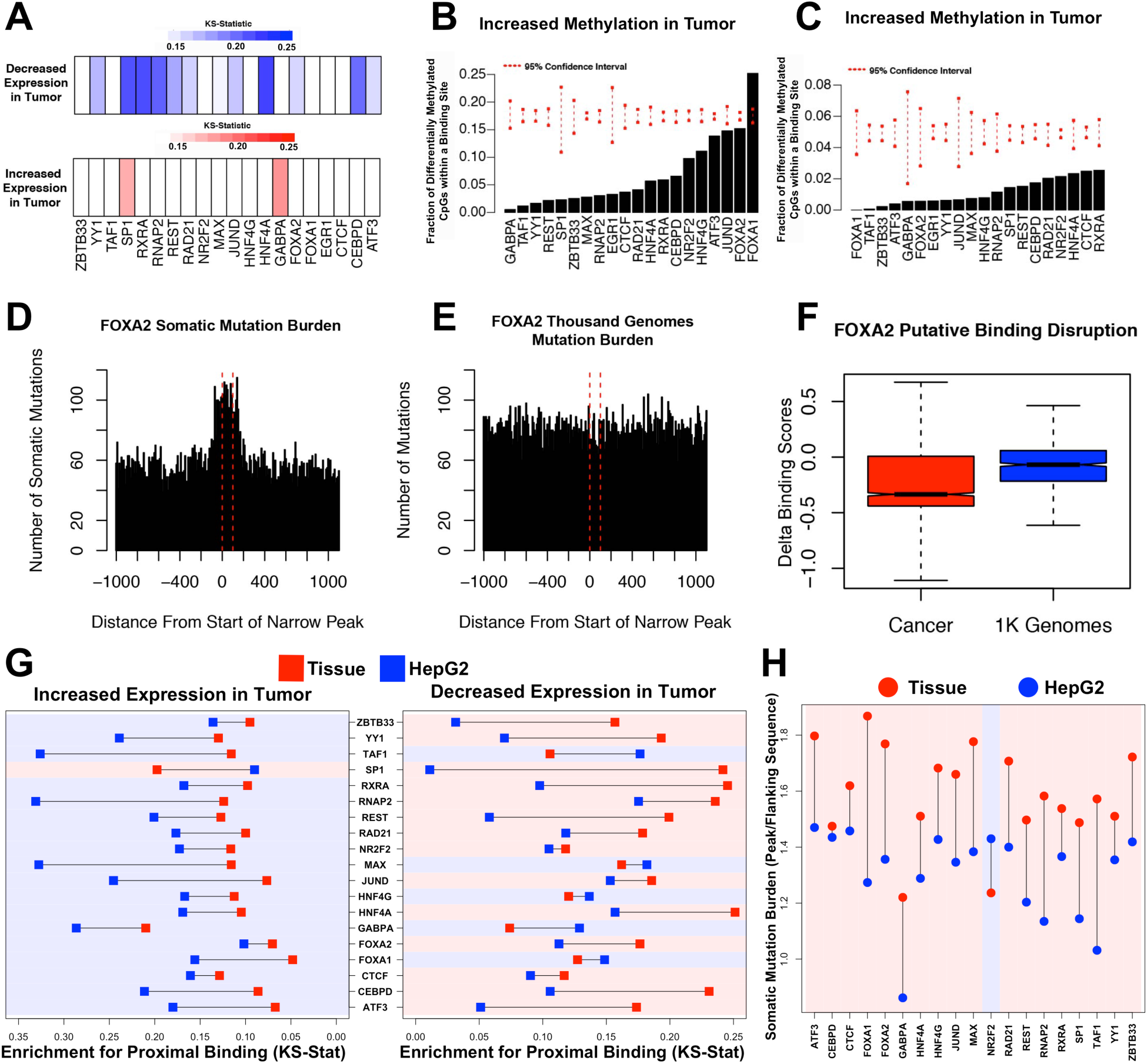
(A) Color bars indicating the KS-test statistics for enrichment of binding of each DAP proximity to the TSS of genes with significantly decreased (blue) or increased (red) expression in tumor tissue compared to adjacent normal tissue. (B and C) Percent of probes with significantly increased or decreased methylation in tumor compared to adjacent normal tissue overlapping a binding site of each DAP. Red dashed lines indicate 95% confidence intervals based on random sampling of an equivalent number of null probes. (D and E) Bars representing the number of somatic mutations (D) or matched 1000 Genome (E) mutations observed at contiguous 10bp bins covering all FOXA2 peaks and flanking 1kb regions. (F) Delta binding scores of all binding site overlapping cancer somatic SNVs (red) or matched 1000 Genome mutations (blue). (G) KS-test statistics for enrichment of binding of each DAP proximal to the TSS of genes with significantly increased or decreased expression in tumor tissue compared to adjacent normal tissue in HepG2 cells (blue) and adult liver tissue (red) for all 19 commonly assayed DAPs. Regions are shaded according to whether HepG2- or tissue-derived peaks display greater enrichment. (H) Mean somatic mutation burden of DAP binding sites over flanking regions in HepG2 cells (blue) and adult liver tissue (red) for all 19 commonly assayed DAPs. Regions are shaded according to whether HepG2- or tissue-derived peaks display greater enrichment.

An analysis of genome-wide DNA methylation at DAP binding sites showed a relative depletion in significant differences between tumor and adjacent normal tissue (Figures 6B and 6C), suggesting that the CpG methylation status at most DAP binding events is unrelated to tumorigenesis. FOXA1, which is known to be methylation-sensitive (Zhu et al. 2016; Bartke et al. 2010), was the only TF whose binding site exhibited a significant enrichment for differential methylation. Nearly 25% of FOXA1 binding sites overlapping CpG dinucleotides had significantly increased methylation in tumor compared to adjacent normal tissue (FDR<0.05). Both *FOXA1* and *FOXA2* genes were expressed at significantly higher levels in TCGA tumor tissue compared to adjacent normal (FDR<0.01), alluding to a functional role for these FOX proteins in hepatocellular carcinoma.

We assessed the degree of somatic variation at all DAP binding sites. Analysis of whole genome somatic single nucleotide variation (SNV) data from 258 hepatocellular carcinoma patients obtained from the International Cancer Genome Consortium (ICGC, http://icgc.org) revealed dramatically increased somatic mutation burden in DAP binding sites compared to flanking regions for the majority of tested DAPs (Figure 6D, Supplemental Figure 17). This effect was not observed in an equivalent sample of variants from the 1000 Genomes Project matched for reference and alternate base pair composition (TG, http://www.internationalgenome.org) (Figure 6E). However, a significant proportion of this increased mutation burden is likely driven by increased tumor mutation rates at cytosine and guanine nucleotides (Supplemental Figure 18), which are enriched in DAP binding sites (Kaiser et al. 2016). FOXA1 and FOXA2 binding sites were outliers with the strongest increased somatic mutation burden after correcting for base-pair composition (*P*=0.01, 0.03), but the robustness of this observation is unclear given that these effects did not survive correction for the scope of hypothesis testing (FDR=0.285, Figure 6D, Supplemental Figure 17, Supplemental Table 15). Interestingly, liver cancer mutations falling within FOXA2 binding sites had a more damaging delta binding score, on average, compared to binding sites overlapping TG SNVs (Wilcoxon *P*=6.4×10^−38^, Figure 6F).

Our tissue-based DAP binding exhibited increased enrichment for proximal binding near genes down-regulated in liver cancer compared to adjacent normal tissue (Paired Wilcoxon *P*=4.6×10^−3^), and conversely HepG2-derived DAP binding sites showed greater enrichment for proximal binding near up-regulated genes (Figure 6G, Paired Wilcoxon *P*=2.1×10^−4^). Moreover, HepG2- derived binding sites exhibited a lower somatic mutation burden relative to flanking regions for all factors except for NR2F2 (Figure 6H, Paired Wilcoxon *P*=3.8×10^−5^). Overall, these data demonstrate that DAP binding sites harbor an increased number of somatic mutations compared to flanking regions, and although a majority of this trend can be attributed to nucleotide composition-related tumor mutational processes, disruption of DAP binding sites may be an important mechanism for altering normal liver gene expression programs. Furthermore, non-cancerous tissue-based ChIP-seq assays may provide insight to regulatory regions disrupted in cancer.

## Discussion

We provide a comprehensive evaluation of DAP binding dynamics by generating 80 independent ChIP-seq datasets comprised of replicate ChIP-seq experiments for 20 different DAPs in liver tissues from two individuals. These data reveal a high level of concordance between both tissues with roughly 75% of DAP binding sites shared between donors. It also highlights interactions between DAPs, with >7% of genes containing more than 6 DAP binding sites within 1kb of their TSS. We demonstrate the degree of DAP co-occupancy is associated with several metrics of activity including conservation, histone marks of open chromatin, and neighboring gene expression. Moreover, DAPs exhibited highly correlated biases in allele occupancy suggesting factors are highly interactive and effects of disruptive sequence variation tend to be shared by neighboring factors. Our data also suggest that DAP influence on neighboring gene expression is highly dependent upon proximity to a gene’s TSS. Both the degree to which allele bias in DAP occupancy is correlated with allele bias in gene expression and the degree to which the number of number of factors bound influences gene expression drops precipitously the further factors are bound from a TSS, suggesting that a substantial majority of DAP activity, at least for the factors assayed, is confined to proximal promoter regions.

Our analysis highlights the value in performing tissue-based ChIP-seq analysis to characterize tissue-specific gene regulation. The DNA-binding proteins we analyzed show a high degree of promoter-proximal binding near genes uniquely expressed in the liver and these DAP binding events are also preferentially enriched in liver specific-eQTL SNPs compared to eQTLs specific to other tissues. These tissue specific correlations were diminished in data generated for the same DAPs in the HepG2 cell line, demonstrating the potential utility of our data in providing a better understanding of the regulation of gene expression programs relevant for in vivo liver physiology, as well as the disruptions that may occur in hepatic disease states. In particular, we find our data to be highly complementary to HepG2 data in terms of understanding cancer-related disruptions in regulatory regions of the genome. While HepG2-derived DAP binding sites showed a higher affinity for proximal binding near genes over expressed in liver tumors compared to adjacent normal tissue, the tissue-derived DAP binding sites we identified here showed a corresponding affinity for proximal binding in genes with decreased expression in tumors. Furthermore, tissue-derived DAP binding sites exhibited higher enrichment for somatic mutations compared to flanking regions than that observed in HepG2 cells. These observations suggest that our data are well suited to inform investigations into cancer-mediated disruptions of cis-regulatory elements in healthy liver. A higher enrichment for overlap with SNPs associated with liver related phenotypes via previous GWAS studies further suggests this utility may expand beyond cancer as well.

We have also provided one possible approach to effectively using our DAP occupancy data to prioritize impactful non-coding sequence variations, which was validated by observations of conservation, allele-specific bias in DAP occupancy at sites of heterozygous SNPs, and *in vitro* reporter assays. Several putative DAP-disruptive eQTL SNPs were associated with relevant phenotypes to liver tissue, including glucose homeostasis, drug metabolism and circulating lipid levels, and therefore represent a promising resource for future mechanistic follow-up experimentation. Of particular interest is the SNP rs11870935, which has previously been identified as GWAS SNP significantly associated with several risk factors for cardiovascular disease including LDL cholesterol and circulating triglyceride levels (Teslovich 2010) and characterized as an intronic/promoter liver eQTL SNP for *KPNB1*, a gene encoding an importin beta subunit critical for nucleocytoplasmic transport regulating cholesterol biosynthesis and insulin resistance via *SREBP* and *NF-kB* respectively (Nagoshi and Yoneda 2001; Wang et al. 2015). We found the alternate allele for this SNP ranked in the top 0.1% of all eQTL SNPs for inducing cryptic *RXRA* binding and was capable of driving increased expression in an *in vitro* reporter assay in HepG2 cells.

There are important limitations to our study. First, we prioritized the breadth of factors assayed, which constrained us to conducting assays on only two individuals. This limits our ability to make conclusions in several important areas such as constructing reasonable estimates of the natural variability of DAP occupancy and identifying robust associations between DAP occupancy and donor demographics like age, sex or ethnicity. Furthermore, we have assayed only a small portion of the known DAPs expressed in humans (Fulton et al. 2009) and repressive factors are particularly underrepresented in our sample. It is likely sampling a larger number of DAPs will provide more insight into the magnitude and complexity of DAP-interactions and uncover more putatively disruptive regulatory sequence variants.

Despite the significant amount of work to be done in fully characterizing the regulatory landscape of the human genome, the application of genomic techniques has shed light on the high level of coordination required for the precise, spatiotemporal control of gene expression. Although painstaking efforts by large consortia have greatly contributed to our understanding of these intricate molecular processes, one notable hurdle that remains is validating functional genomic data generated in cell culture models in tissues. To this end, we generated this comprehensive dataset of genome-wide DAP binding events in two healthy human liver tissues and provided an analytical overview providing insight into gene regulation in tissue and disease related disruptions. We believe that this work will serve as an important resource to the research community and will further facilitate a broader functional genomic investigation of DAP functions across an array of additional human tissues. By systematic integration of genomic datasets from complementary efforts using cellular and tissue models, the complexity of regulatory genome architecture will be steadily uncovered.

## Methods

### ChIP-seq experimentation and DAP interaction analysis

#### Tissue procurement

Liver tissue was obtained from both deceased donors and flash frozen with liquid nitrogen at the time of organ procurement through Mid-America Transplant Services/Washington University (St. Louis, MO). Research consents from donor families were obtained.

#### Chromatin immunoprecipitation and next-generation sequencing

ChIP-seq experimentation was performed using a previously established method (Savic et al. 2013). Briefly, for each ChIP-seq replicate in each liver, an independent dry pulverization was performed in Covaris tissueTUBES attached to glass vials. After pulverization, tissue was collected in the attached vial, fixed, washed, and stored as a pellet at -80^°^C. To avoid bias from batch effects, each replicate for each ChIP experiment was prepped together in a single batch (resulting in two total batches) and the ChIP assay and subsequent library preparations were conducted as previously reported (Reddy et al. 2012). Antibodies used for ChIP-seq assays are listed in Supplemental Table 16. All antibodies have been previously used in conjunction with cell line based ChIP experiments conducted at HudsonAlpha and made publically available through the ENCODE Project. All ChIP-seq libraries were run on an Illumina HiSeq2500 sequencer using 50bp single-end sequencing. Binding sites were identified using the MACS peak caller using an mfold cutoff of 15 (Zhang et al. 2008) while enriched binding motifs were identified through MEME (Bailey et al. 2009). Narrow peaks were defined as 100bp segments of DNA centered on the peak summit. ChIP-seq replicates were used to identify concordant peaks for downstream analysis. Concordant peaks were defined as narrow peaks that were present in both replicate and overlapped by at least one base pair.

#### DAP-interaction analysis

Normalized binding intensity at the union of all narrow peak DAP binding sites in each tissue was obtained by merging all narrow peaks, determining the total number of reads that were mapped to each merged region for each ChIP experiment, normalizing these values by the total number of reads mapped for each ChIP experiment (in other words, converted to reads per million), and constructing a Spearman correlation matrix for all factors in each tissue (Quinlan 2014). To identify DAP binding clusters, hierarchical clustering was performed on the correlation matrices for all pair-wise factor combinations and plotted as a heatmap. The correlation matrix was also used to construct a network diagram of DAP interactions with the R package “qgraph” (Epskamp et al. 2012). Histone modifications were obtained from the Epigenome Roadmap Project (Consortium et al. 2015). The total amount of genome covered by merged narrow peaks from the adult liver was compared to an equivalent number of matched null sequences generated from the Beer Lab galaxy (http://kmersvm.beerlab.org) “Generate Null Sequence” function allowing 2% repeat and GC content error. To assess conservation at binding sites with increasing numbers of DAPs bound, narrow peaks were first merged in a manner that tracked the number of overlapping factors bound at a given site. Next, mean GERP scores were determined for each site using base-wise GERP rejected substitution scores from the Sidow lab website (http://mendel.stanford.edu/SidowLab/downloads/gerp/hg19.GERPscores.tar.gz). To obtain histone modification intensities at each merged peak, BigWig files were obtained for adult liver tissue from the Epigenome Roadmap data portal (http://egg2.wustl.edu/roadmap/webportal/) and converted to BEDgraph files with the UCSC genome browser (https://genome.ucsc.edu/goldenpath/help/bigWig.html) “bigWigtoBedGraph” function. Mean BedGraph intensities, calculated as fold change over a reverse cross-linked control, were calculated for each merged peak. Each DAPs relative overlap with a histone mark was calculated by obtaining liver tissue narrow peak BED files for each histone modification from the data portal and calculating the fraction of DAP narrow peaks that overlapped with a given histone modification peak.

#### Whole-genome sequencing and allele-specific binding analysis

10X Chromium, whole genome sequencing and phased BAM and VCF files were generated from frozen liver tissue from each donor via the 10X genomics longranger pipeline by the HudsonAlpha Genomic Services Lab (https://gsl.hudsonalpha.org/information/10X). Allele bias in the adult liver was assessed with ChIP-seq data using the R package “BaalChIP” (DeSantiago et al. 2016). BaalChIP requires binary alignment/map (BAM) files generated from replicate ChIP-seq experiments, a BAM file from genomic DNA (gDNA) sequenced, and a VCF file with heterozygous SNPs overlapping previously called ChIP-seq binding sites. BaalChIP calculates expected allele frequencies from gDNA BAM files, filters SNPs with “MAPQ” value <15 and “QUAL”<10, filters SNPs in regions with UCSC mappability scores<1 or present in UCSC blacklisted mappability tracks and repeat regions, adjusts for reference mapping bias, filters possible homozygous SNPs, and uses a beta-binomial Bayesian model to detect allele specific binding events. BaalChIP output consists of corrected allelic ratios of reads overlapping each heterozygous SNP and a Boolean variable indicating whether significant allele bias was detected at SNP. A pair-wise Spearman correlation matrix was generated comparing allele bias at SNPs passing BaalChIP quality filters for each pair of factors. At least 25 peaks had to pass quality filters for a pair of factors to be included in the analysis (30 out of 380 possible pairs were removed).

### Regulation of liver-specific gene expression

#### RNA extraction and sequencing and alignment

Four independent tissue pulverization, as described previously, were performed on each liver. RNA was extracted from pulverized samples using Qiagen RLT buffer +1% BME. RNA was purified from 350 uL of tissue lysate using the Norgen Animal Tissue RNA purification kit. RNA-seq libraries were generated using Tn-RNA-seq, a transposase-mediated construction method, as previously described (Gertz et al. 2012b). All replicates were prepped in a single batch and and sequenced using an Illumina HiSeq 2500 to generate 50bp paired-end reads. Sequencing reads were aligned using a previously described pipeline (Alonso et al. 2017). Briefly, reads were trimmed using TrimGalore with default settings prior to being aligned and converted into a raw count table using STAR (Dobin et al. 2012). Percent unique alignment was consistent across samples and ranged from 75 to 81% uniquely aligned reads. Transcripts were aligned to hg19 reference genome and the genomic coordinates of all transcripts were obtained from the Ensembl genome browser (http://useast.ensembl.org/index.html) grch37_E75 gene transfer format (GTF) file. Gene expression levels were normalized to transcript length as well as total read depth, and expressed as transcripts per kilobase million (TPM).

#### Allele-specific expression analysis

Allele specific expression was calculated for each expressed heterozygous SNP using the GATK “ASEReadCounter” function according to previously described best practices (Castel et al. 2015). SNPs with a read depth less than 10, less than 5 reads assigned to each allele, had a “MAPQ” value <10 and “QUAL” value <2, and had a UCSC genome browser mappability score <1 were filtered prior to analysis. A simple binomial test using the R “binom.test” was performed to assess the significance of the observed number of reference reads out of the total number of reads at each SNP with the null ratio assumed to be the average genome-wide reference bias of 0.5397 after performing quality filtering. Only SNPs present within the same phase set (labeled PS within the phased .VCF file INFO column) were used in DAP-binding and expression allele bias correlations. Ninety-five percent confidence intervals for this analysis were generated by randomly shuffling DAP overlapping and expression SNP pairs that were within a given distance threshold and assessing the correlation 100 times.

#### Gene set-ChIP binding proximity analysis

Reactome pathway information was obtained from http://www.reactome.org/pages/download-data/ (Fabregat et al. 2016). Promoter-proximal pathway enrichment was calculated as follows. For a given pathway, the distribution of distances from the TSS of each gene to the nearest binding site for a given factor was compared to the distribution of TSS to nearest binding site distances for the entire transcriptome as previously described (Savic et al. 2016) and significance was determined using the non-parametric Kologomorov-Smirnov test. Concordant narrow peaks that were present in all replicates from both livers were used for pathway analysis. Mean pathway expression was calculated as the mean TPM normalized expression for each gene in a given pathway. Median RPKM normalized expression levels of each transcript for each tissue were obtained from the GTEx data portal (GTEx_Analysis_v6p_RNA-seq_RNA-SeQCv1.1.8_gene_median_rpkm.gct.gz). Tissue specific transcripts were defined as transcripts that had an RPKM value greater than 2 and a 5-fold higher expression in a given tissue compared to the average expression in all remaining GTEx tissues. Replicate BAM files for each protein factor analyzed in our human tissue donors, except for EGR1, were obtained from previous work in our group in HepG2, which is publically available at the ENCODE data portal (https://www.encodeproiect.org). BAM files were processed as described above for the liver tissue ChIP-seq experiments prior to analysis.

#### DAP-binding site overlap with GTEX eQTL SNPs

GTEx eQTL sites were obtained with permission from the GTEx project download portal “GTEx_Analysis_v6p_eQTL.tar” file. Binding site-eQTL overlaps were assessed with adult liver replicate concordant narrow peaks. Enrichment for liver eQTL overlap was assessed by comparing the observed significant (GTEx Q-value <0.05) eQTL overlap to 1000 random permutations of non-significant SNPs (GTEX Q-value>0.05) that passed the GTEx consortium quality filters. Null SNPs were matched to significant eQTL SNPs for distance to a TSS and minor allele frequency based on the Thousand Genomes Project data (http://www.internationalgenome.org). Matching was performed by binning SNPs into twenty quantiles based on distance to the nearest TSS and then separately binning SNPS into twenty quantiles based minor allele frequencies. Successive rounds of randomly sampling was then performed such that number of SNPs sampled from each bin was equivalent to the number of significant eQTLs present in each bin at each round. Enrichment for overlap with liver-specific eQTLs was performed in a similar manner except percent overlap with eQTLs with a GTEx q-value < 0.05 only in liver was compared to tissue specific eQTLs in all other tissue types. The number of tissue-specific eQTLs ranged from ~2500 to ~25000 and liver fell roughly at the median with ~5000 eQTLs. To assess relative eQTL enrichment between DAPs a linear model was fit regressing the number of observed significant eQTL overlaps by the total number of binding sites for each factor (Number of Significant eQTL Overlapping Sites ~ Number of Total Sites) with the R “lm” function. The relative enrichment for each factor was assessed as the residual error for each factor from the expected overlap fit by the linear model. This enrichment was compared to the coefficient of variation or relative standard deviation (σ/μ) to all genes with TSSs within 5Kb (and a variety of other thresholds) of a given factor’s binding site.

#### DAP-binding site overlap with NHLBI GRASP GWAS SNPs

The NHLBI GRASP2.0 catalog of GWAS studies (https://s3.amazonaws.com/NHLBI_Public/GRASP/GraspFullDataset2.zip) was used to obtain genomic coordinates of published GWAS SNPs. All SNPs present in the catalog were used for analysis. Prior to analysis, phenotype terms relevant to liver physiology were selected for comparing HEPG2 and Tissue ChIP-seq experiments. Next, overlapping binding sites from all DAPs (excluding EGR1, which wasn’t assayed in HepG2) were merged into a master binding site list and GWAS SNPs associated with each phenotype in the GRASP catalog located in a HEPG2 or adult tissue binding site were counted. A simple Fisher’s exact test was performed comparing the proportion of each individual GWAS phenotype term’s SNPs that fell within a binding site to the total number GWAS SNPs in the GRASP catalog that overlapped a binding site.

#### Prioritizing impactful non-coding variation

##### Support vector machine training

Support vector machines (SVMs) were trained on replicate-concordant narrow peak sites from the adult liver using a method previously established (Lee 2016; Ghandi et al. 2014). Briefly, genome sequence was obtained in FASTA format for each narrow peak using the Bedtools “getfasta” command. A GC content, repeat content, and length matched set of null peak sites 10 times greater in number than the number of concordant peak sites observed for each factor was obtained from the kmersvm galaxy web site (http://kmersvm.beerlab.org) using the “Generate Null Sequence” function. SVMs were trained on narrow peak and matched background sequences using gapped 10-bp kmers and allowing for 3 non-informative bases using the “gkmtrain” executable obtained from the ls-gkm github webpage (https://github.com/Dongwon-Lee/lsgkm). All other settings were left at default. This resulted in 20 SVMs, one for each DAP analyzed. Model performance was determined using receiver-operator characteristic area under the curve (ROC-AUC) and precision recall area under the curve (PR-AUC) on predictions from 5-fold cross validation. ROC-AUC and PR-AUC curves were constructed using the “PRROC” package in R.

##### Delta binding score and GERP Correlation

To determine the relationship of conservation and the importance of each base pair in predicting binding site status in the SVM, we performed individual *in silico* mutations of every reference sequence nucleotide in each narrow peak site by mutating them to the three other possible alternates for each DAP. This resulted in 100bp sequences with a single base-pair change for each base in the reference narrow peak, generating a total of 300 mutant sequences for each narrow peak. Each mutant sequence was scored with the original SVM trained on its respective DAP. The classifier value obtained for the 3 possible alternate alleles at each reference base was averaged and then subtracted from the reference sequence classifier value to obtain a delta binding score for each base within each narrow peak, indicative of the relative importance of each base for SVM classification performance. These scores were correlated with GERP-RS conservation and allele-specific bias at each site.

##### Scoring GTEx eQTL SNPs

To identify GTEx liver eQTL SNPs likely to disrupt DAP binding or cause *de novo* DAP binding, we obtained a 100-bps of genome sequence centered on each liver eQTL SNP using the BEDTools “getfasta” command to generate two, 100bp sequence windows containing the reference or the alternate allele. The reference and alternate sequences were subsequently scored with the each of the 20 SVMs trained on each DAP analyzed. The reference classifier value was subtracted from the alternate allele to obtain a delta binding score as described above. Putative eQTL SNPs with DAP loss of binding had to overlap with a replicate-concordant narrow peak binding site, have a delta enhancer score in the bottom 0.1 percentile of all eQTL SNPs and possess a mutant classifier value less than 0. Conversely, putative eQTL SNPs with DAP gain of binding were defined as those not overlapping with a replicate-concordant narrow peak site and were required to yield a delta enhancer score in the top 99.9 percentile and possess a mutant classifier value greater than 0. Putative gain and loss of binding eQTL SNPs were queried in the NHLBI GRASP2.0 catalog of GWAS studies for potential disease relevance (Eicher et al. 2015). Enrichment for GRASP phenotype terms was assessed by Fisher’s exact test comparing the proportion of eQTL SNPs scoring in the top 1% of delta binding scores and significantly associated with a given GRASP phenotype term to the proportion of SNPs significantly associated with a given GRASP phenotype term in the entire population of liver eQTL SNPs.

##### In vitro reporter assays

All reporter assays were performed in HepG2 hepatocarcinoma cell lines. We randomly selected five liver eQTL SNPs in the top 0.1% of all delta binding scores. Reference and mutant sequences for each SNP were cloned into the pGL4.23 vector (Promega) multiple cloning site upstream of a minimal promoter driving luciferase (*luc2*) expression using Gibson Assembly (Gibson Assembly Master Mix, NEB). Plasmid DNA was extracted from three separate colonies with the Spin Miniprep Kit (Qiagen) and sequence verified with Sanger sequencing (MCLAB, San Francisco, CA). Each colony was treated as a separate biological replicate for a given sequence. HepG2 cells were seeded at 40,000 cells per well in antibiotic free DMEM with 10% FBS in a 96-well plate. After 24 hours, 300ng of plasmid DNA for each biological replicate was transfected into HepG2 cells using FuGENE (Promega) in duplicate, resulting in 6 total replicates (3 biological X 2 technical) per reference or mutant sequence. Luciferase activity was measured 48-hours post-transfection with a 2-second integration time on a LMax II 384 Luminometer (Molecular Devices). Background subtracted luminescence values for each SNP were z-scored. Significance in expression was determined using a 2-tailed Student’s T-test.

#### Liver cancer analyses

##### RNA-seq analysis

RNA-seq, DNA methylation and copy number variation data was obtained with permission from NCI-GDC data portal (https://gdc-portal.nci.nih.gov/). Raw RNA-seq read counts were obtained from matched tumor and adjacent normal pairs for 49 TCGA patients. Differential expression between tumor and adjacent normal was determined using DESeq2 (Love et al. 2014) using default settings. Enrichment for proximal binding near differentially expressed genes was performed using the Kologomorov-Smirnov test-based approach described above by comparing differentially expressed genes (DESeq2 FDR<0.001, n=9,832) with a background set of genes that received at least one sequencing read in one sample.

##### DNA methylation analysis

Methylation beta values from the Illumina Infinium HumanMethylation450 chip array were obtained for 50 matched tumor and adjacent normal tissue pairs. Probes with missing values (n=105,836) or with variance less than 0.001 (n=91,495) were removed prior to the analysis. Differential methylation was determined using the R package “samr” (Tusher et al. 2001). The non-parametric “SAM” function was used with the “resp.type” set to “Two class paired” and “nperms” set to 3,000. Differentially methylated probes were defined as those with a median beta value difference greater than 0.1 between tumor and adjacent normal tissues and a SAM q-value less than 5 (n=85,088). The percent of replicate-concordant narrow peaks overlapping a differentially methylated probe for each DAP was determined using the BEDTools “intersect” function. This was compared to the percent in overlap using 1000 randomly-selected probes of equivalent number.

##### Somatic copy number variation analysis

Somatic Affymetrix SNP 6.0 based DNA copy number data was obtained for 376 tumors from the TCGA data portal. Only deep deletions and amplifications classified as genome segments with a normal-masked segment mean with an absolute value greater than 1.0 were used for analysis. The percentage of deleted or amplified binding sites for each DAP in at least 5 patients was computed using the Bedtools “intersect” command on CNV data obtained for all patients.

##### Somatic single nucleotide variant analysis

Somatic mutation data was obtained from the International Cancer Genome Consortium (ICGC) data release 23 from the LIRI-JP project consisting of somatic SNVs from 258 patients with liver cancer (Fujimoto et al. 2016). A total of 2,691,076 SNVs were found in this dataset. Somatic mutation burden at the binding site and flanking regions for each DAP were obtained by intersecting the somatic mutation coordinates with contiguous 10-bp bins spanning the binding site regions, along with 1000bp of flanking sequence. A similar analysis was performed on the Thousand Genomes (TG) project SNVs by randomly sampling mutations from the Phase 3 variant call format file while conserving the nucleotide mutation proportions relative to the observed cancer mutations (Auton et al. 2015). A simple logistic regression was performed with the “glm” function in R to test for significant enrichment of somatic mutation burden within binding sites for each DAP compared to flanking regions, while correcting for GC content as follows:

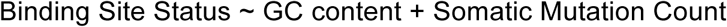

where binding site status is a binary variable indicating whether a 10-bp window is contained within a narrow peak or a flanking region, GC content is a continuous variable ranging from 0 to 1 indicating the fraction of G or C nucleotides in a 10bp window and Somatic Mutation Count is the number of SNVs found within a 10bp bin. To identify putative binding-disruptive somatic SNVs, a similar method was used as described above for GTEx eQTL SNPs. Briefly a 100-bp of sequence centered on the SNV was obtained and scored with all 20 SVMs trained on each DAP for the reference and alternate allele. The delta-binding scores were calculated as the difference in SVM classifier or fitted decision values between the mutant and wild-type sequence.

#### Data Access

All data is freely available at the ENCODE data portal (https://www.encodeproiect.org) under the sample accession numbers ENCDO882MMZ and ENCDO060AAA.

## Acknowledgments

We thank Christopher Partridge for his assistance designing and conducting plasmid reporter assays, Krysta Engel and Mark Mackiewicz for general advice on the manuscript, and Mike Snyder for sharing liver tissue for analysis. We also thank the HudsonAlpha Genomics Services Laboratory, especially Dr. Shawn Levy and Ms. Angela Jones, for their contributions to the large-scale DNA sequencing performed for the ChIP-seq, RNA-seq, and whole genome sequencing experiments in this study. This work was supported by the National Institute of Health (NIH) grants U54 HG006998-0 and 5T32GM008361-21 (RCR). SJC is supported by the HudsonAlpha Tie the Ribbons Fund, the UAB CCTS (NIH 1UL1TR001417-01), and support from the State of Alabama.

## Author Contributions

DS, RM, SC, and GC conceived of the study

DS and KN prepped samples for RNA-seq and ChIP-seq

RR and DS performed computational analysis

AH and RR performed reporter assays

RR, DS, GC, AH, SC, and RM contributed to writing the manuscript

## Disclosure Declarations

The authors declare no conflicts of interest.

## Supplemental Figure Legends

**Supplemental Figure 1.** Correlation network diagram showing relationships between normalized DAP binding intensities at all observed binding sites of both livers. Blue and red nodes indicate adult and child binding sites, respectively, and the width of each edge indicates the strength of the spearman correlation between each factor.

**Supplemental Figure 2.** Boxplots showing the spearman correlations of normalized binding intensities between the same DAP across donors and all DAP pairs for the adult and child donor.

**Supplemental Figure 3.** Bar plot indicating the proportion of factors bound at merged DAP binding sites in which FOXA1 is bound (blue) or is absent (red).

**Supplemental Figure 4.** (A) Mean GERP score of genomic regions containing an increasing number of factors bound. (B-E) Epigenome Roadmap project histone mark fold enrichments over background at sites with increasing numbers of factors bound.

**Supplemental Figure 5.** (A) Pie chart representing the percent of genes containing a specified amount of bound DAPs within 1Kb of their TSS for the child liver. Color scale reflects the number of neighboring DAPs required to be included in each slice. (B) Expression level of genes binned by the number of DAPs bound within 1Kb of their TSS for the child liver.

**Supplemental Figure 6.** Scatter plot indicating the number of RNA-seq reads mapping to the reference and alternate allele at expressed heterozygous SNPs in the adult liver. Red dots indicate SNPs that hold a Bonferonni corrected P-value<0.05 for differential allele expression while blue dots indicate SNPs with a P-value>0.05.

**Supplemental Figure 7.** Plot indicating the correlation in DAP binding allelic bias and neighboring gene expression allelic bias at a variety of binding site-TSS distance thresholds for each DAP. A positive correlation indicates a DAP tends to predominantly bind on the same allele that is predominantly expressed. A correlation of zero is indicated by the blue dashed line. Ninety five percent confidence (red) intervals calculated by randomly shuffling all SNP pairs that met a distance threshold 100 times and computing a null correlation distribution.

**Supplemental Figure 8.** Diagram visually demonstrating the TSS-DAP binding proximity-based pathway analysis for identifying cis-regulatory networks for two DAPs.

**Supplemental Figure 9.** Dots represent KS-statistic of enrichment for proximal binding of each DAP to liver (red/purple), skin (blue) and cortex (green) -specific genes in child liver tissue.

**Supplemental Figure 10.** Enrichment for proximal binding to liver specific expression analysis, as shown in Figure 3C-D, over a range of fold enrichment threshold cutoffs. From top to bottom, tissue specific transcripts were defined as those with a mean RPKM of 2 in a given tissue of interest and 2, 3, 4, and 5-fold increased mean expression in the tissue of interest relative to the mean expression across all other tissues. Dots represent KS-statistic of enrichment for proximal binding of each DAP to liver (red/purple), skin (blue) and cortex (green) -specific genes in adult liver tissue (left) and HepG2 cell line (right).

**Supplemental Figure 11.** Red dot indicates the percent of liver-specific eQTLs overlapping binding sites for each DAP relative to the boxplots representing the percent overlap of all other GTEx tissue-specific eQTLs.

**Supplemental Figure 12.** (A) Red dot indicates the number of eQTLs falling within a HepG2 DAP binding site relative to the gray boxplots representating 1000 randomly sampled null SNPs matched for distance to TSS and minor allele frequency. Plot is the HepG2 compliment to Figure 3A. (B) Red dot indicates the percent of liver-specific eQTLs overlapping binding sites for each HepG2 DAP relative to the boxplots representing the percent overlap of all other GTEx tissue-specific eQTLs. Plot is the HEPG2 compliment to Figure 3B.

**Supplemental Figure 13.** Mean GERP score of all adult peaks for each DAP.

**Supplemental Figure 14.** (A) ROC curve AUC values for SVMs trained on each DAP. (B) Precision-recall curve AUC vales for SVMs trained on each DAP. Representative plot for SVM models in ATF3. Plots the frequency of each 10mer in ATF3 binding sites against its SVM weight. Color of each point reflects the GC content of the 10mer.

**Supplemental Figure 15.** Paneled pairs of plots for each DAP overlapping at least one SNP with significant allele bias and at least one SNP with a delta binding score falling in the top percentile (15/20). The left figure for each factor is a scatterplot demonstrating the correlation between delta binding score and the observed ChIP-seq allele bias at heterozygous SNPs overlapping a binding site. Red points indicate SNPs with significant bias towards an alternate allele, blue points indicate bias towards the reference allele, and black points indicate no significant allele bias was observed. The right figure consists of boxplots of binding site-overlapping heterozygous SNPs predicted to be in the top ~1% for decreasing binding affinity (red), to be in the top ~1% for increasing binding affinity (blue), and to have no significant impact on binding affinity (black). Y-axis indicates the fraction of ChIP reads mapping to the reference allele.

**Supplemental Figure 16.** Fisher’s p-value of enrichment for NIH GRASP GWAS catalog terms associated with putative DAP-disruptive SNPs relative to background enrichment in all GTEx liver eQTL SNPs.

**Supplemental Figure 17.** Panels for each DAP showing the number of cancer somatic mutations (top left) or matched 1000 Genome (top right) observed in each DAP peak and flanking 1Kb region. The percent base pair composition at each position (bottom left) and delta binding scores for cancer somatic mutations and matched 1000 Genome mutations (bottom right) are plotted as well.

**Supplemental Figure 18.** The number of liver cancer somatic mutations observed for each nucleotide per occurrence of that nucleotide in the genome. Colors within each bar represent the proportion of nucleotides to which the reference allele is mutated.

